# Genetic architecture of trophic adaptations in cichlid fishes

**DOI:** 10.1101/2022.06.03.494688

**Authors:** Leah DeLorenzo, Victoria DeBrock, Aldo Carmona Baez, Patrick J. Ciccotto, Erin N. Peterson, Clare Stull, Natalie B. Roberts, Reade B. Roberts, Kara E. Powder

## Abstract

Since Darwin, biologists have sought to understand the evolution and origins of phenotypic adaptations. The skull is particularly diverse due to intense natural selection such as feeding biomechanics. We investigate the genetic and molecular origins of trophic adaptation using Lake Malawi cichlids, which have undergone an exemplary evolutionary radiation. We analyze morphological differences in the lateral and ventral head among an insectivore that eats by suction feeding, an obligate biting herbivore, and their F_2_ hybrids. We identify variation in a series of morphologies including mandible width, mandible length, and buccal length that directly affect feeding kinematics and function. Using quantitative trait loci (QTL) mapping, we find that many genes of small effects influence these craniofacial adaptations. Intervals for some traits are enriched in genes related to potassium transport and sensory systems, the latter suggesting correlation between feeding structures and sensory adaptations for foraging. Craniofacial phenotypes largely map to distinct genetic intervals, and morphologies in the head do not correlate. Together, these suggest that craniofacial traits are mostly inherited as separate modules, which confers a high potential for the evolution of morphological diversity. Though these traits are not restricted by genetic pleiotropy, functional demands of feeding and sensory structures likely introduce constraints on variation. In all, we provide insights into the quantitative genetic basis of trophic adaptation, identify mechanisms that influence the direction of morphological evolution, and provide molecular inroads to craniofacial variation.

## INTRODUCTION

Understanding the patterns and origins of variation is a key challenge within both developmental biology and evolutionary biology. A structure with significant morphological diversity is the skull, with variation across and within many clades of vertebrates including fishes [1-3], birds [4-6], reptiles [7, 8], and mammals [9-13]. A critical selective pressure faced by craniofacial structures is trophic niche specialization, with skull morphology directly feeding into biomechanical performance and fitness [14]. These forces shape a complex geometry of the skull, with morphological variation deriving from the cumulative effects of genetics, developmental processes, environmental effects, and functional interactions [15-20].

An iconic system for morphological variation is cichlid fishes, which have undergone one of the most rapid diversifications in vertebrates [21, 22]. A hallmark of their adaptive radiation is the diversity of craniofacial structures, which are intimately connected to their feeding niche and ecology [2, 23, 24]. Cichlids, like other teleost fishes, have evolved multiple disparate feeding strategies including suction feeding, biting, and ram feeding, each of which is associated with a suite of phenotypic adaptations [24]. Despite this range of craniofacial morphologies in cichlids, a major ecomorphological axis of variation in cichlids distinguishes two of these strategies, suction feeding and biting [2, 25]. On one end of this axis are suction feeders. These animals eat from the water column by generating a high rate of flow into the mouth that overcome any flow in the opposite direction or attempts by mobile prey to swim away [26-29]. Morphologically, this is accomplished through a large buccal cavity and restricted mouth size that confer an ability to generate pressure differentials in the oral cavity [26, 27]. Production of the pressure differential is enhanced through a relatively long mandible that allows quick movements of the jaw [28, 30-33]. Further, large eyes in suction feeders may increase vision to provide an advantage in hunting prey [34], but may also constrain the size of jaw muscles needed for mandible movement [35]. On the alternate end of this morphological spectrum are fishes that feed by scraping/biting attached algae or crushing shelled invertebrates. These fish trade off speed in mandible movements for power with jaw closing, primarily conferred by a shorter lower jaw [28, 30-32].

Cichlids from independent radiations have undergone similar divergences in craniofacial morphology between fish that suction feed versus bite [25, 36, 37], and this trend extends more broadly across fishes as well [38-40]. This pattern suggests that genetic, developmental, or functional constraints are limiting or biasing the direction of morphological evolution in the skull [41-43]. For example, coordinated changes could be driven by “supergene” regions [44-46] or biomechanical demands of ecological niches may cause convergent evolution of form (e.g. [47]). A full understanding of the patterns of morphological variation, as well as the number and effects of genes that underlie these shapes, is necessary to clarify which aspects of head anatomy demonstrate covariation, have increased evolutionary flexibility, or are simpler versus more complex phenotypes.

Here, we use two species of cichlids to investigate the adaptation of craniofacial morphology and the genetic basis of this variation. Both *Labidochromis caeruleus* and *Labeotropheus trewavasae* live in rocky habitats of Lake Malawi, but feed by suction feeding and scraping, respectively [23]. Fishes of the *Labidochromis* genus are typically insectivores that suction feed or pluck their prey from the water column [23]. Alternatively, fishes of the *Labeotropheus* genus strictly feed by biting algae that is attached to rocky substrates [23]. We first quantify a series of morphological adaptations in the lateral and ventral craniofacial skeleton in these species. We then utilize quantitative trait loci (QTL) mapping in a population of *Labidochromis* x *Labeotropheus* F2 hybrids to ask if these traits are controlled by the same genetic intervals or distinct loci, and thus inherited as a module or independently, respectively. Finally, we examine candidate genes and pathways enriched by gene ontology (GO) term analysis to uncover molecular mechanisms that may influence craniofacial morphological diversity. Overall, these data will elucidate genetic factors that influence diversity in trophic adaptations of the craniofacial skeleton and drive major morphological variation in the skull.

## MATERIALS AND METHODS

### Fishes and pedigree

All work was completed under animal protocol 140-101-O approved by the Institutional Animal Care and Use Committee (IACUC) at North Carolina State University. A single *Labidochromis caeruleus* female was crossed with a single *Labeotropheus trewavasae* male to create one F_1_ family, that was subsequently incrossed to produce a hybrid F_2_ population of 447 fishes. Hereafter, *Labidochromis caeruleus* and *Labeotropheus trewavasae* will be referred to as their genus name. Fish were reared in aquaria under standard feeding with flake food for five months, at which time they were euthanized with buffered MS-222 for morphological analysis. Lateral and ventral images of each specimen were taken using an Olympus digital camera under standardized lighting conditions in a lightbox. A color standard and scale were included in each picture.

### Linear measures of head shape variation

Measures were taken on 10 parental specimens per species of *Labidochromis* and *Labeotropheus*, and either 447 F_2_ hybrids for lateral analysis or 319 F_2_ hybrids for ventral analysis. From photographs of the lateral body, we measured standard length (snout to caudal peduncle), head length (snout to opercle), head depth (anterior insertion of the dorsal fin to the insertion of the pelvic fin), length from the snout to the insertion of the pelvic fin, preorbital length (snout to anterior edge of the eye), eye diameter, and mouth angle (Figure 1b). Eye area was calculated from eye diameter measurements. Measures of the ventral anatomy included mandible width, mandible length, width from the posterior of the opercle to midline, length from the posterior of the opercle to the joint of the mandible and palatoquadrate, and mandible angle (Figure 1d). Measurements were taken using ImageJ software as number of pixels and were then converted into centimeters using the scale in each photo. To remove the effects of allometry, all measures were converted to residuals by normalizing to standard length using a data set with both parental species and their hybrids. Further analysis was conducted in R, including ANOVAs, Tukey’s Honest Significant Difference post-hoc tests, and correlations.

**Figure 1.**
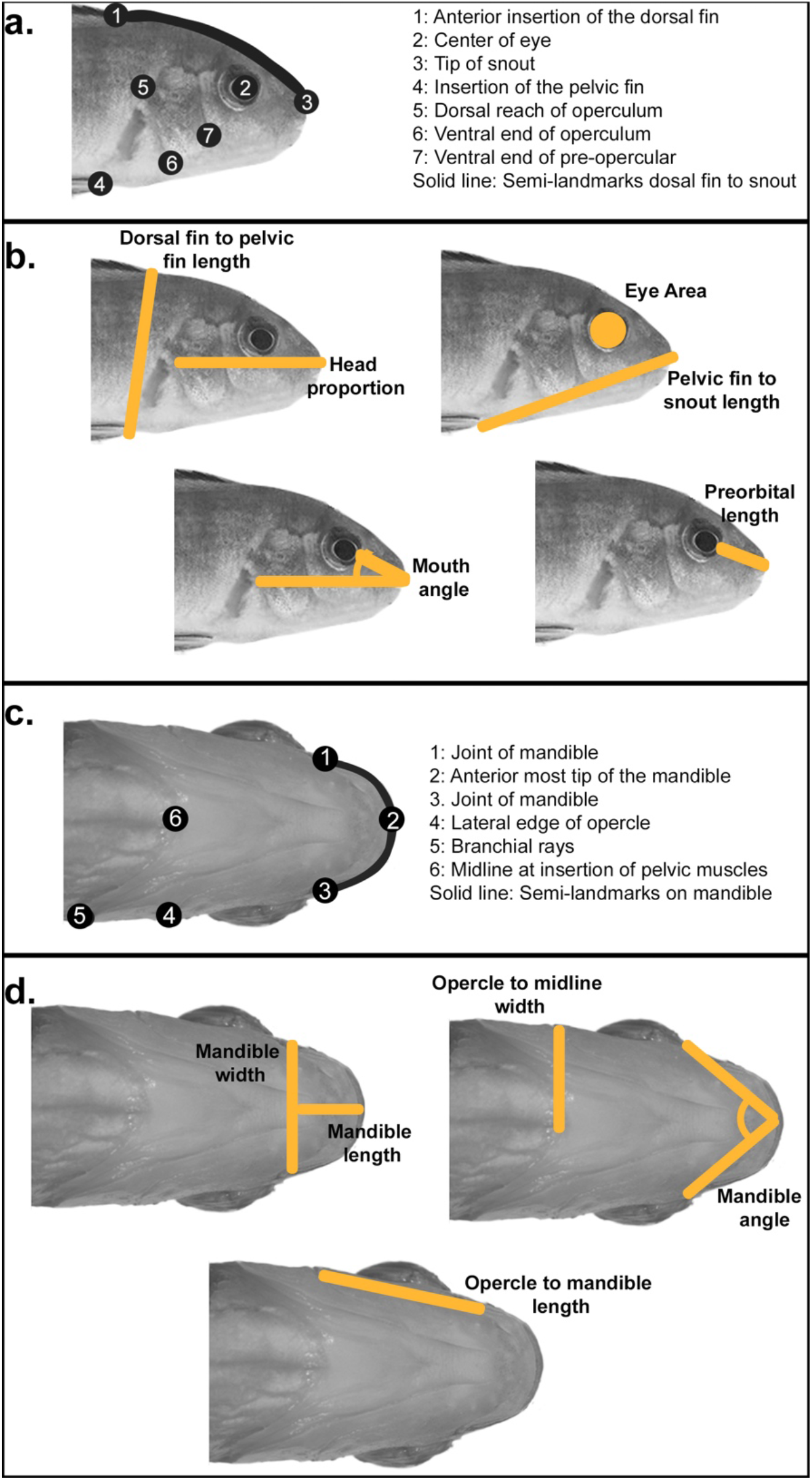
Measures used to assess lateral and ventral head shape. **(a**,**c)** Geometric and **(b, d)** linear measures were used to assess head shape changes with functional implications for feeding biomechanics.

### Geometric morphometric shape analysis

Geometric morphometric shape analysis was used to further quantify head shape variation. A series of homologous landmarks were chosen highlighting lateral and ventral craniofacial anatomy important to feeding mechanics (Figure 1a, Figure 1c). In both cases, we only analyzed one side of the specimen, avoiding the side in which there were body dissections posterior to the pectoral fins. X,Y coordinates of all landmarks were collected and extracted from photos using the tpsDig2 software package [48]. These data were uploaded into the R package geomorph, in which Procrustes superimposition was used to remove variation due to size, rotation, and position of landmarks to leave variation only due to shape. As with the linear data, the effects of allometry were removed through size correction and regression of shape on standard length. All geometric morphometric analyses were conducted on a data set including both parental species and their hybrids.

### Genotyping with ddRAD-sequencing

Genomic DNA was extracted from caudal fin tissue using DNeasy Blood and Tissue kits (Qiagen). RADseq libraries were prepared as previously described [49], including double digestion and indexing, then sequenced on Illumina Hiseq with 100bp paired end reads (North Carolina State University Genomic Sciences Laboratory core facility). The program process_radtags (Stacks, version 2), was used to process raw sequencing data including demultiplexing, truncating reads to 150bp, and filtering of low-quality reads. Processed reads were aligned to the *Maylandia zebra* UMD2a reference genome using BWA with the mem algorithm. The programs pstacks, cstacks, and sstacks (Stacks, version 1) were used to identify and catalogue RAD markers in the parental and F_2_ hybrid samples. Finally, markers with alternative alleles in the parental species were called as AA or BB genotypes using the program genotypes (Stacks, version 1), requiring a minimum stack depth of 3 to export a maker in a specific individual. The A allele was inherited from the *Labidochromis* granddam and the B allele from the *Labeotropheus* grandsire.

### Generation of the linkage map

The genetic map was generated using the package R/qtl [50] and in-house R scripts. Markers were first sorted into linkage groups according to their position in the *M. zebra* UMD2a reference genome. Markers were removed from the data set if they were located on unplaced scaffolds with more than 40% of missing data, or in linkage groups with more than 20% missing data. A chi-square test was performed on the remaining markers using the geno.table function. Those markers with a distorted segregation pattern and a Bonferroni-corrected p-value < 0.01 were discarded from the dataset. The initial map was generated based on estimated pairwise recombination frequencies using est.map and est.rf functions. Markers in linkage groups that were not initially flagged as misplaced were removed if they increased the size of the map by at least 6 centimorgans (cM) and flanking markers were < 3Mb apart. Markers that were in unplaced scaffolds were integrated into a linkage group if they had a recombination frequency < 0.15 with at least 5 markers from that linkage group. Any other markers that were in unplaced scaffolds that did not meet the above criteria were removed. If markers had irregular relationships between their recombination frequency and position in the genetic map, they were rearranged manually to minimize crossover events; these are likely due to being located in structural variants or misassembled sections of the reference genome. Genotyping errors were identified using the function calc.errorlod and set as missing data if they had a LOD score ≥ 3. The linkage map was refined with a non-overlapping window algorithm that selected one marker in a 2cM window with the least amount of missing data. Finally, the function est.map was used to estimate the final map and the maximum likelihood estimate of the genotyping error rate (0.0001). The final map was 1239.5 cM in total size, with 22 linkage groups, 1180 total markers and 42-81 markers per each linkage group.

### Quantitative trait loci (QTL) mapping

We conducted multiple-QTL mapping (MQM) using the R/qtl package [50-52] following [53]. Scripts are described and available in [54]. First, an initial scan for QTL was done using the onescan function in R/qtl [50]. Putative QTL with a LOD approaching or above 2.5 were used to build a more robust statistical model. The MQM method uses these putative QTL as cofactors in follow-up scans and verifies each cofactor by backward elimination. The use of cofactors in the final model aids in the accurate detection of QTL and assessment of their effects [53]. The statistical significance of each QTL was determined using 1000 permutations on the final model. For QTL peaks meeting 5% (significance) or 10% (suggestive) level, 95% confidence intervals were calculated using Bayes analysis. Details of QTL mapping including cofactors used in the model, significance levels, confidence intervals, and allelic effects are in Table S3.

### Candidate gene annotation and enrichment analysis

The markers are named based on contig and nucleotide positions in the *M. zebra* reference genome, M_zebra_UMD2a assembly. Gene symbols, ID, and chromosomal positions for candidate genes in each QTL interval were retrieved from the NCBI genome data viewer (https://www.ncbi.nlm.nih.gov/genome/gdv) gene track for *M. zebra* annotation release 104. If the upper and lower limits of a QTL interval were mapped to unplaced scaffolds, the closest marker that mapped to a placed scaffold was used to determine candidate gene information. Gene names for each candidate were retrieved using the NCBI gene ID and the Database for Visualization and Integrated Discovery (DAVID) [55].

Gene ontology (GO) term enrichment analysis was performed with the functional annotation tool in the Database for Visualization and Integrated Discovery (DAVID) [55, 56]. NCBI gene ID (entrez gene ID) for candidate genes in QTL intervals were used as a query. Analysis was run for each individual trait, pooling multiple QTL as applicable, as well as bulk analysis of all lateral QTL and all ventral QTL. A p-value of 0.05 with a Fishers exact probability test was used to denote significance for terms in GO analysis.

## RESULTS AND DISCUSSION

### Lateral head shape variation

Lateral skull shape is distinct between parental species *Labidochromis* and *Labeotropheus* for all linear measures (Figure 2a-f and Table S1). Their F_2_ hybrids are largely intermediate in phenotype, though in some cases such as length of the preorbital region (Figure 2d) surpass the range of the parental species. *Labidochromis* fish have an overall longer and deeper head than *Labeotropheus* given a similar body size. Specifically, *Labidochromis* compared to *Labeotropheus* parentals have an increased proportion of the body that is the head (p<1e-7, Figure 2a), a longer distance between the dorsal fin and pelvic fin (p<1e-7, Figure 2b), and larger eye (p<1e-7, Figure 2e). Further, the mouth of *Labidochromis* fish is angled towards the front, rather than towards the ventral side of the body as in *Labeotropheus* (p<1e-7, Figure 2f). Finally, *Labidochromis* showed an increased length between the snout and pelvic fin (p=2.2e-6, Figure 2c). Coupled with a more modest, though still significant, enlargement of the preorbital region (p=0.018, Figure 2d), this suggests that the opercular region of these fishes is also distinct.

**Figure 2.**
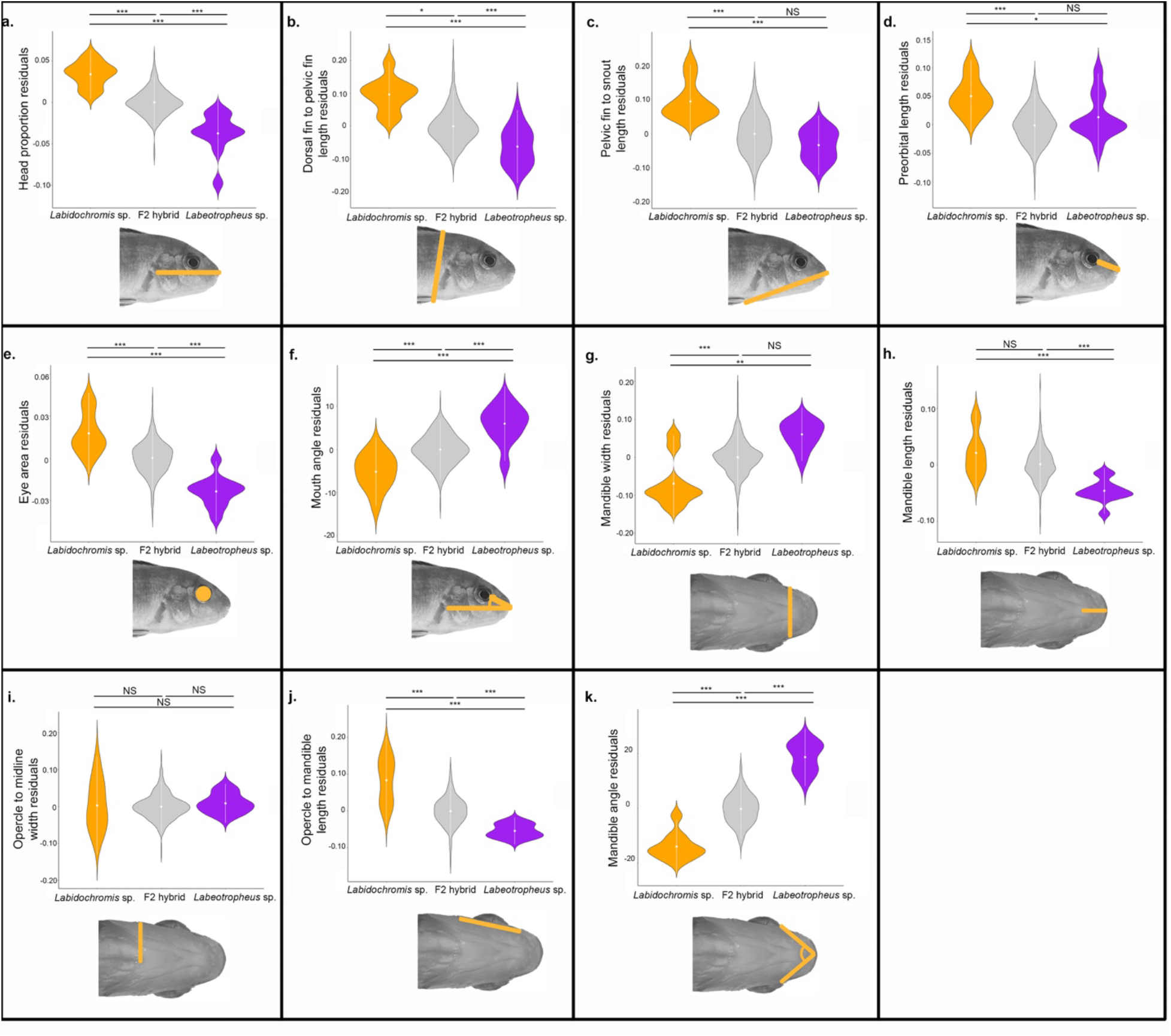
Phenotypic differences among *Labidochromis* sp., *Labeotropheus* sp., and their F2 hybrids. Phenotypes measured are indicated by illustration and include **(a)** head proportion, measured as head length/standard length, **(b)** dorsal to pelvic fin length, **(c)** snout to pelvic fin length, **(d)** length of the preorbital region of the head, **(e)** eye area, **(f)** mouth angle, **(g)** mandible width, **(h)** mandible length, **(i)** opercle to midline width, **(j)** length from the opercle to the mandible, and **(k)** angle formed from posterior ends of the mandible to the midline. Significance in violin plots is based on ANOVA analysis followed by Tukeys HSD (data in Table S1; p-values indicated by * <0.05, ** <0.01, *** <0.005, NS >0.05).

Geometric morphometrics provided more detailed insights into shape differences, including within the opercular region of the head. The first five principal components (PCs) described (75.2% total shape variation [TSV]) in lateral shape in *Labidochromis* sp., *Labeotropheus* sp., and their F2 hybrids (Figure 3a, Figure 3c, and Figure S1). PC1 lateral (22.4% TSV) differentiated the two parental species (p<1e-7), with *Labidochromis* species associated with a positive PC1 lateral score that describes a longer head with a more posterior eye placement (Figure 3a and Figure S1b). Based on linear measures, this shift in eye position is due to both a larger preorbital region (Figure 2d) and a larger eye area (Figure 2e). As suggested by linear measures, PC1 lateral shape differences show that *Labidochromis* has a larger opercular region, while the operculum in *Labeotropheus* only extends about halfway between the eye and insertion of the pelvic fin (Figure 3c). PC2 lateral, PC3 lateral, and PC4 lateral were not significantly different between the parentals (p=0.071, p=0.99, and p=0.77, respectively, Table S1) and thus represent shape variation largely present in the F_2_ hybrids. PC2 lateral (17.2% TSV) predominantly described the relative length of the head, with a negative PC2 lateral score characterizing head anatomy that has a longer profile from snout to dorsal fin and a pelvic fin that is inserted closer to the opercle (Figure S1c). PC3 lateral (13.3% TSV) depicted coordinated changes in both head length and depth, with a negative score representing a deep, short head with a steep craniofacial profile and reduced opercular region (Figure S1d). Notably, a steep craniofacial profile in cichlids has been associated with an ability for the skull to withstand increased biting forces [57]. PC4 lateral (11.4% TSV) describes differences in the dorsal-ventral depth of the opercular region, as well as the dorsal-ventral positioning of the eye (Figure S1e). Finally, PC5 lateral (11.0% TSV) distinguishes the two parental species (p=0.012). *Labidochromis* parentals are associated with a more negative PC5 lateral score and a reduced opercle bone (Figure S1f).

**Figure 3.**
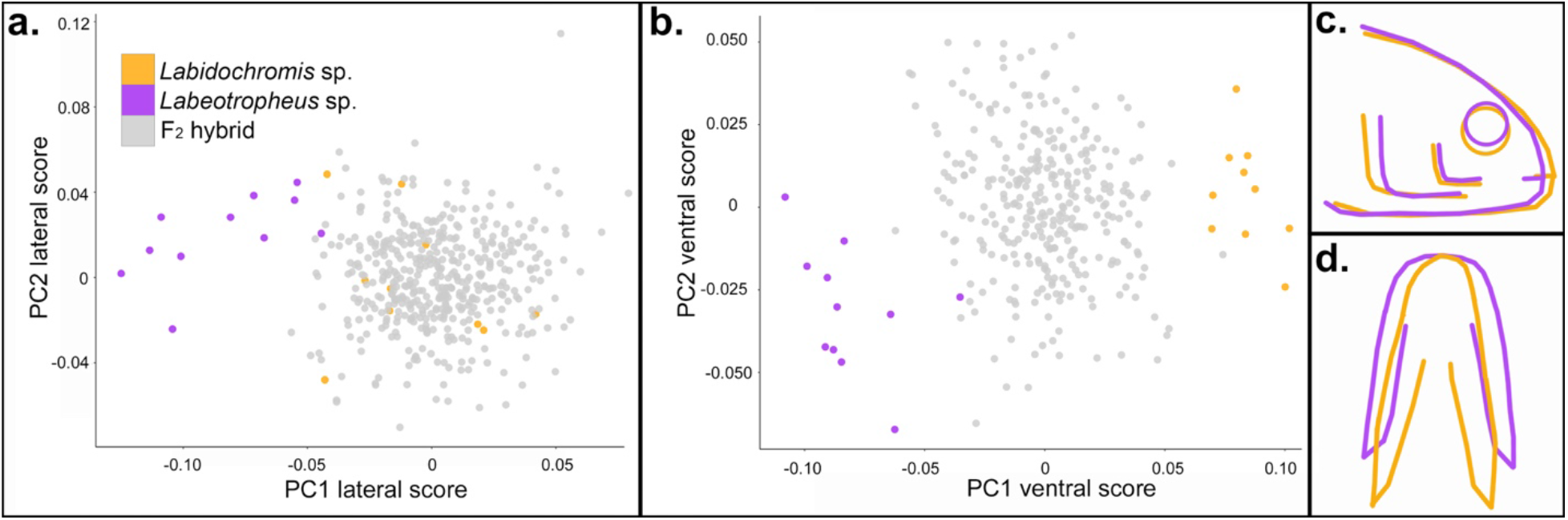
Geometric morphometric phenotypes among parentals and hybrids. Multivariate analysis of shape quantifies differences in overall morphology in the **(a, c)** lateral and **(b, d)** ventral anatomy. Shapes described by each principal component are described in the text and visualized in Figure S1. Average shape **(c, d)** of *Labidochromis* sp. (orange) and *Labeotropheus* sp. (purple) based on **(a**,**b)** highlights phenotypic variation between alternate feeding strategies.

### Ventral head shape variation

Compared to *Labeotropheus, Labidochromis* parental fish have a decreased mandible width (p<1e-7, Figure 2g), increased mandible length (p=4e-7, Figure 2h), and longer length of the opercular region (p<1e-7, Figure 2j). Mandible angle assesses the relative proportions of the lower jaw, with an increased measure indicating increased width, decreased length, or both, in the case of *Labeotropheus* (p<1e-7 compared to *Labidochromis*, Figure 2k). These shape changes combine with a similar width at the opercle (p=0.93), Figure 2i), the only measure that was not distinct between parentals. This results in a more triangular ventral shape for *Labidochromis* and a more rectangular shape for *Labeotropheus* parentals (Figure 3d).

Relative mandible length and width also dominated geometric morphometric analysis of the ventral skeleton. The first three ventral principal components cumulatively describe 76.2% TSV in ventral craniofacial anatomy. PC1 describes 43.6% TSV, with the parental species defining the extremes (p<1e-7, Figure 3b). *Labidochromis* parents are associated with a positive PC1 ventral score and a narrower, arched mandible versus the wide and flat mandible shape of *Labeotropheus* (Figure 3b, Figure 3d, and Figure S1g). PC2 ventral (18.7% TSV) is also distinct between parentals (p=8.2e-5, Figure 3b), with a narrow mandible, increased distance of the opercular region, and pectoral fin musculature shifted to the anterior (Figure S1h). PC3 ventral (13.9% TSV) describes relative mandible length without an accompanying change in the width (Figure S1i) and is not significantly different between *Labidochromis* and *Labeotropheus* parentals (p=0.31).

Combining both lateral and ventral shape variation demonstrates the multiple ways *Labidochromis* and *Labeotropheus* have craniofacial biomechanics that are adapted to their feeding niches. *Labidochromis* sp. pluck or suction feed insects within Lake Malawi [23]. Their longer mandibles (Figure 2h) allow more velocity transmission during jaw movement [32], critical for capture of mobile prey. This is combined with a narrow mandible (Figure 2g) that opens into a longer and wider opercular and buccal region (Figure 2i-j), forming a triangular ventral shape (Figure 3d). The large expansion possible in the buccal cavity of *Labidochromis* causes high velocity and acceleration of water flowing into the mouth, containing the invertebrate prey; this water flow is increased by a narrow mouth opening (Figure 2g, Figure 3d) [26, 58, 59]. On the other hand, *Labeotropheus* sp. are herbivorous grazers that scrape or shear immobile algae from rocks or other substrate using their mandible [23]. The short mandible (Figure 2h) of *Labeotropheus* represents a tradeoff of speed of jaw movement for high transmission of force with jaw closing [32]. This is combined with a downturned mouth (Figure 2f) and a short, wide, and flat preorbital and mandibular region (Figure 2d, Figure 2g, Figure 3c, and Figure 3d). Together, these are thought to enhance foraging efficiency for *Labeotropheus* by providing a large oral area and structures that are used as a fulcrum to leverage attached algae from their substrate [23].

### Genetic basis of body shape

Quantitative trait loci (QTL) mapping was used to assess the genetic architecture that underlie these adaptive morphologies. Mapping of all 19 traits (11 lateral and 8 ventral measures) including both linear (Figure 2) and geometric measures (Figure 3 and Figure S1) of shape identified 23 genetic intervals that contribute to phenotypic differences in head shape in *Labidochromis* x *Labeotropheus* hybrids (Figure 4, Figure S2, Figure S3, and Table S3). Between one and three QTL mapped to 12 of the 22 linkage groups. These QTL explained 3.3-7.0% of the total variation for each trait (Figure S3 and Table S3), indicating that each of these traits is controlled by many genes of small effects. Even for the trait with the most QTL, PC2 lateral shape, the 5 QTL combine to explain only 23.8% of the total coordinated variation (Figure S3) in head length, craniofacial profile, and pelvic fin insertion (Figure S1c). The allelic effects within this QTL (Figure S3) further suggest a complex genetic architecture, with the allele inherited from the *Labidochromis* parent contributing to a higher PC2 lateral score for QTL on LG7 and LG10, the *Labeotropheus* allele associated with a higher value for the QTL on LG2, and heterozygous animals having the largest PC2 lateral score for the QTL on LG6 and LG23. Given that cichlid species continue to segregate and exchange a set of ancestral polymorphisms [60-64], this genetic variation is all likely to contribute to craniofacial divergence and feeding adaptation within the cichlid flock.

**Figure 4.**
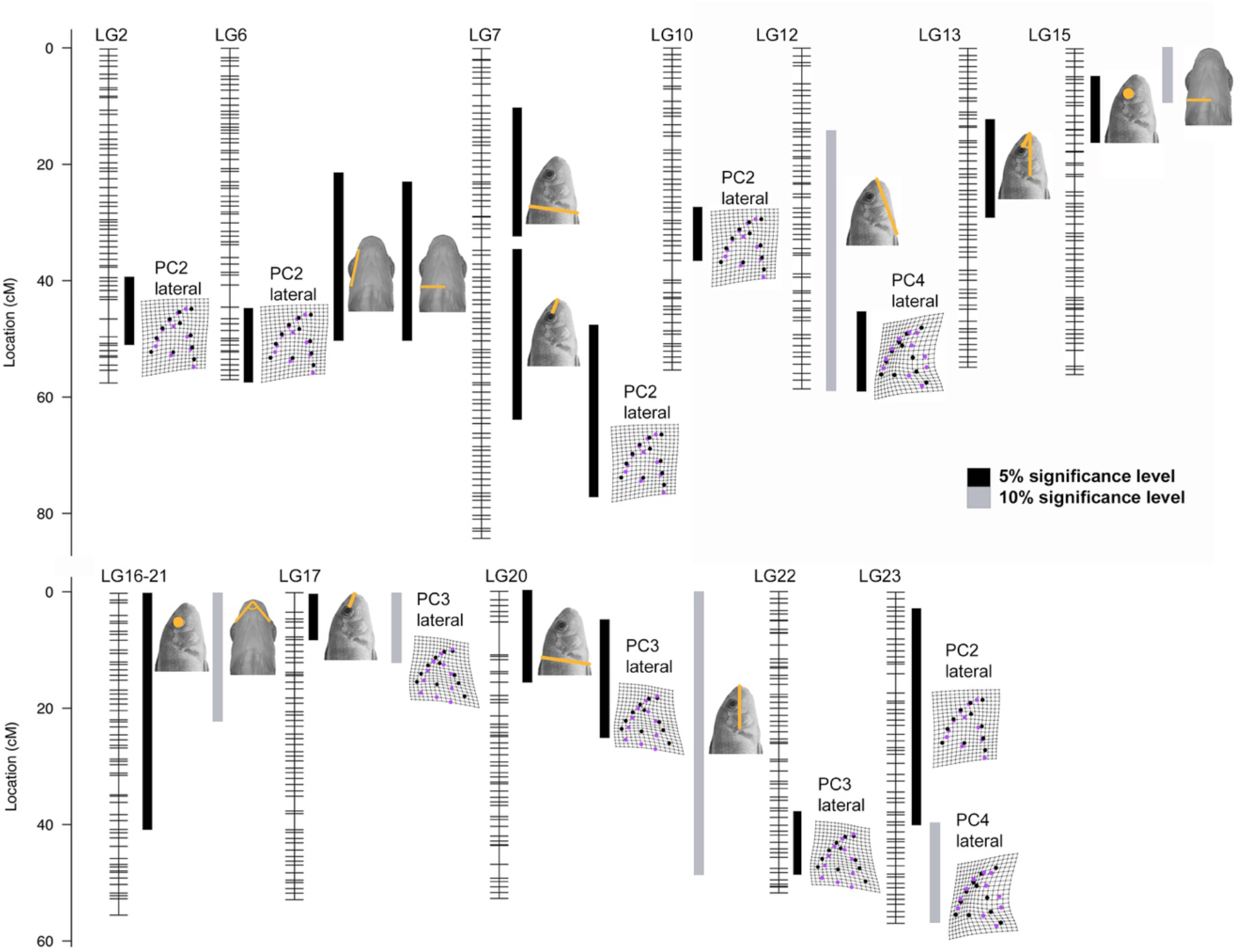
Quantitative trait loci (QTL) mapping identifies 23 intervals associated with head shape variation in hybrids of *Labidochromis* and *Labeotropheus*. Each linkage group (LG, i.e. chromosome) is indicated with genetic markers noted by hash marks. The phenotype related to each QTL region is indicated by illustrations. Black bars are significant at the 5% genome-wide level, while gray bars are suggestive, meeting the 10% genome-wide level. Bar widths indicate 95% confidence interval for the QTL, as calculated by Bayes analysis. QTL scans at the genome and linkage group level are in Figures S2 and S3. Details of the QTL scan including markers and physical locations defining each region are in Table S3.

While QTL were distributed across linkage groups, seven linkage groups had overlapping QTL intervals (Figure 4 and Table S3). Four of these overlapping regions included a linear measure and a principal component from geometric morphometrics, where the principal component also includes variation in that linear measure. For instance, there are three overlapping QTL intervals on LG20 which describe relative head length, depth of the head from the dorsal fin to the pelvic fin, and PC3 lateral shape (Figure 4). PC3 lateral shape includes major variation in the anterior-posterior length and dorsal-ventral depth of the head (Figure S1d), explaining why these phenotypes map to a common genetic interval. Likewise, preorbital length varies in both PC2 lateral and PC3 lateral shape (Figures S1c and Figure S1d). QTL for the preorbital region overlap with QTL for PC2 lateral and PC3 lateral on LG7 and LG17, respectively (Figure 4 and Table S3). Finally, the length of the pelvic fin insertion point to the tip of the snout is part of PC4 lateral shape (Figure S1e), and QTL for these traits overlap on LG12 (Figure 4 and Table S3).

Overlap of QTL may also lead to a coordinated change in shape. However, aside from effects of allometry (i.e. correlation with standard length, Table S2), no phenotypes showed morphological correlation (0.8 < r < -0.8) with each other in the F2 hybrids. Correlations of phenotypes ranged from -0.65 to 0.78 with a mean of 0.027 (Table S2). This suggests that the morphological traits are largely inherited as modular units rather than as a set of coordinated phenotypes. Despite this, we noted linkage groups that have overlapping QTL for both lateral and ventral shape variation. LG6 contains a QTL cluster for PC2 lateral shape, opercle to mandible length, and opercle to midline ventral width (Figure 4 and Table S3). Genetic intervals associated with eye area overlap with opercle to midline width on LG15 and mandible angle on LG16-21 (Figure 4 and Table S3); for all these QTL, the allele inherited from *Labidochromis* increases each of these measurements (Figure S3, Table S3). This common genetic basis, and even sometimes common allelic effects, indicate that a single gene or linked genes in this interval may have pleiotropic effects on feeding adaptations. However, the fact that phenotypes were largely controlled by distinct QTL and showed minimal correlations (Table S2) means that distinct feeding morphologies could theoretically evolve independently and recombine into new patterns. This modular pattern would increase the morphological variability possible in cichlids (i.e. be more evolvable) [65-69]. Despite this, three independent, large-scale radiations of cichlids in the African Rift-Lakes have generated animals with similar trophic specializations that share remarkable similarities in their craniofacial morphologies [25, 36, 37]. Thus, despite largely being independent in terms of genetic structure, morphological disparity is constrained. Our data suggests this is predominantly due to functional demands of feeding and strong natural selection on feeding performance, rather than a genetic constraint [70-72].

### Gene Ontology (GO) analysis

More work is needed to narrow down and determine the specific effects of candidate genes within QTL intervals (Table S4), but GO analysis was used as a start to identifying trends and pathways that are enriched. Members of the Wnt signaling pathway were significantly enriched (p=0.046, Table S5) for mouth angle, though we note this is only a single QTL on LG13. There is a strong relationship between the mouth angle and the steepness of the craniofacial profile (see solid line in Figure 1a), with a shallow profile leading to a narrow mouth angle and jaw facing forward. Alternatively, a steep profile is associated with *Labeotropheus* sp. [73], an increased mouth angle (Figure 2f) and ventrally angled jaws. Wnt signaling plays a pivotal role in shape of the craniofacial profile, with increased Wnt signaling causing a retention of larval phenotypes and a steep facial profile in cichlids [73, 74]. Based on the function of Wnt signaling in craniofacial development across vertebrates, this is likely through alteration of cellular proliferation and outgrowth [4, 73, 75-77] and precocious bone deposition [73, 78, 79].

Four traits are statistically significant for changes in potassium transport: head proportion (p=0.018), the distance between the dorsal and pelvic fins (p=0.018), PC2 lateral shape (p=0.031), and PC4 lateral shape (p=0.024) (Table S5). This common signal for head proportion and dorsal to pelvic fin length is likely driven by the fact that these traits have an overlapping QTL on LG20. Further, both PC2 lateral (Figure S1c) and PC4 lateral (Figure S1e) include variation in both of these linear measures. Potassium could have numerous influences on craniofacial morphology as this mineral regulates cell proliferation [80], chondrogenesis [81], osteoclast [82] and osteoblast [81, 83] differentiation, and bone mineralization [81, 83]. Potassium can also influence pathways critical for facial and bone development such as Bmp signaling [75, 81, 84], which is also associated with mandibular adaptation in cichlids [32]. Finally, mutation of potassium channels can lead to a series of developmental syndromes that include craniofacial morphologies that mimic evolved variation in cichlids. For example, Andersen-Tawil syndrome is characterized by a broad facial width and mandibular hypoplasia [85-87], while Birk-Barel syndrome results in a narrow forehead, micrognathia, and cleft or high-arched palate [86, 88] (see Figures 2g-i).

It is perhaps unsurprising that eye area was enriched for the GO terms olfaction and sensory transduction (p=1.25e-6 and p=5.5e-5, respectively, Table S5), given the common developmental origin of sensory structures [89, 90]. However, both mandible angle and a combined analysis of all ventral skeletal morphologies were also enriched for genes associated with these terms (p=3.1e-4 to p=2.26e-8, Table S5). This may be due to coordinated adaptations for feeding strategies as olfaction and sight are important for identifying mobile prey prior to suction feeding [34, 91, 92]. However, this may also be due to functional and spatial constraints, wherein a narrow face or large jaw musculature restricts the space available to develop large eyes [35].

## CONCLUSIONS

Craniofacial variation is prodigious across vertebrates, with direct impact on feeding strategy and fitness. Here, we identify the genetic basis for a series of adaptations related to suction feeding versus biting, including overall head proportions, mandible shape, ventral width, and dimensions of the buccal cavity. These phenotypes are not correlated and largely share independent genetic architecture. Our data thus suggests that craniofacial morphologies are likely constrained due to functional demands rather than similar genetics.

## SUPPLEMENTARY MATERIALS

Supplementary data includes additional geometric morphometric details, QTL scans and details summarized in Figure 4, and tables with statistical analyses of phenotypes, QTL scan details, candidate genes in QTL intervals, and GO analysis.

## ACKNOLEDGEMENTS AND FUNDING

This work was supported by NSF CAREER IOS-1942178 (KEP), NIH P20GM121342 (KEP), NIH R15DE029945 (KEP), NSF IOS-1456765 (RBR), and an Arnold and Mabel Beckman Institute Young Investigator Award (RBR).

## AUTHOR CONTRIBUTIONS

KEP and RBR conceptualized the research. ACB, ECM, PJC, and NBR performed animal husbandry, photography, and collections. NBR prepared sequencing libraries. KEP, LD, and VD performed phenotypic measurements. KEP, LD, VD, ECM, ACB, and RBR analyzed data. KEP and LD wrote the initial paper with edits and review from all authors. KEP and RBR administered the project and acquired funding.

## DATA AVAILABILITY

Data is accessible at Dryad [link to be provided prior to publication]. These files include phenotypic measures, TPS files for geometric morphometric analysis, and genotypes used for quantitative trait loci mapping.

