## Supplementary Material for "Genetic architecture of trophic adaptations in cichlid fishes"

Supplemental table legends and figures with legends for DeLorenzo et al.

1. Legends for Supplemental tables, provided as separate .xlsx files (pg. 1-2)
    a. Table S1. Statistical significance for phenotypic measures.
    b. Table S2. Correlations between all pairs of phenotypic traits in *Labidochromis* x *Labeotropheus* $F_2$ hybrids.
    c. Table S3. Details of quantitative trait loci (QTL).
    d. Table S4. Candidate genes for head shape variation QTL
    e. Table S5. Gene ontology (GO) analysis for enriched processes in candidate genes for QTL.
2. Figure S1. Shapes indicated by PC scores following geometric morphometric analysis. (pg. 3)
3. Figure S2. Genome-wide QTL scans. (pg. 4-5)
4. Figure S3. QTL scans and allelic effects for QTL. (pg. 6-17)

**Table S1. Statistical significance for phenotypic measures.** Significance among each pair of *Labidochromis* parents, *Labeotropheus* parents, and their $F_2$ hybrids were assessed by ANOVA and Tukeys HSD on size-corrected, residual values. These data support significance notations in Figure 2.

**Table S2. Correlations between all pairs of phenotypic traits in *Labidochromis* x *Labeotropheus* $F_2$ hybrids.** Calculations are for $F_2$ hybrids only. Except for correlations with standard length, all measures are size corrected. PC scores from geometric morphometric analysis were size corrected prior to assessing correlation with standard length. Values that are >0.8 or <-0.8 are in bold.

**Table S3. Details of quantitative trait loci (QTL).** For each QTL, we list cofactors used to generate models. Markers names and genetic positions in centimorgans (cM) are noted for the peak of the QTL and the upper and lower bounds of the 95% confidence interval. Marker names include the physical location on the linkage group, with names referring to the contig and nucleotide position in the *M. zebra* UMD2a assembly. LOD values, percent phenotypic variance explained by that QTL, allelic effects, and additive, dominance, and heritability calculations are for the peak marker in the QTL. QTL listed in gray text are suggestive at the 10% significance level, while those in black text meet 5% genome-wide significance based on values indicated.

**Table S4. Candidate genes for head shape variation QTL.** For the indicated 95% confidence interval, we include candidate gene names in the *M. zebra* annotation release 104, NCBI gene ID number, and physical positions in the *M. zebra* genome UMD2a assembly.

**Table S5. Gene ontology (GO) analysis for enriched processes in candidate genes for QTL.** Over-represented biological process GO terms (p<0.05) associated with genes in QTL intervals for each individual trait, all lateral QTL, and all ventral QTL. Each GO term includes the number of genes and NCBI Gene ID that are in the QTL interval(s) and match the category, fold enrichment, and p-value.

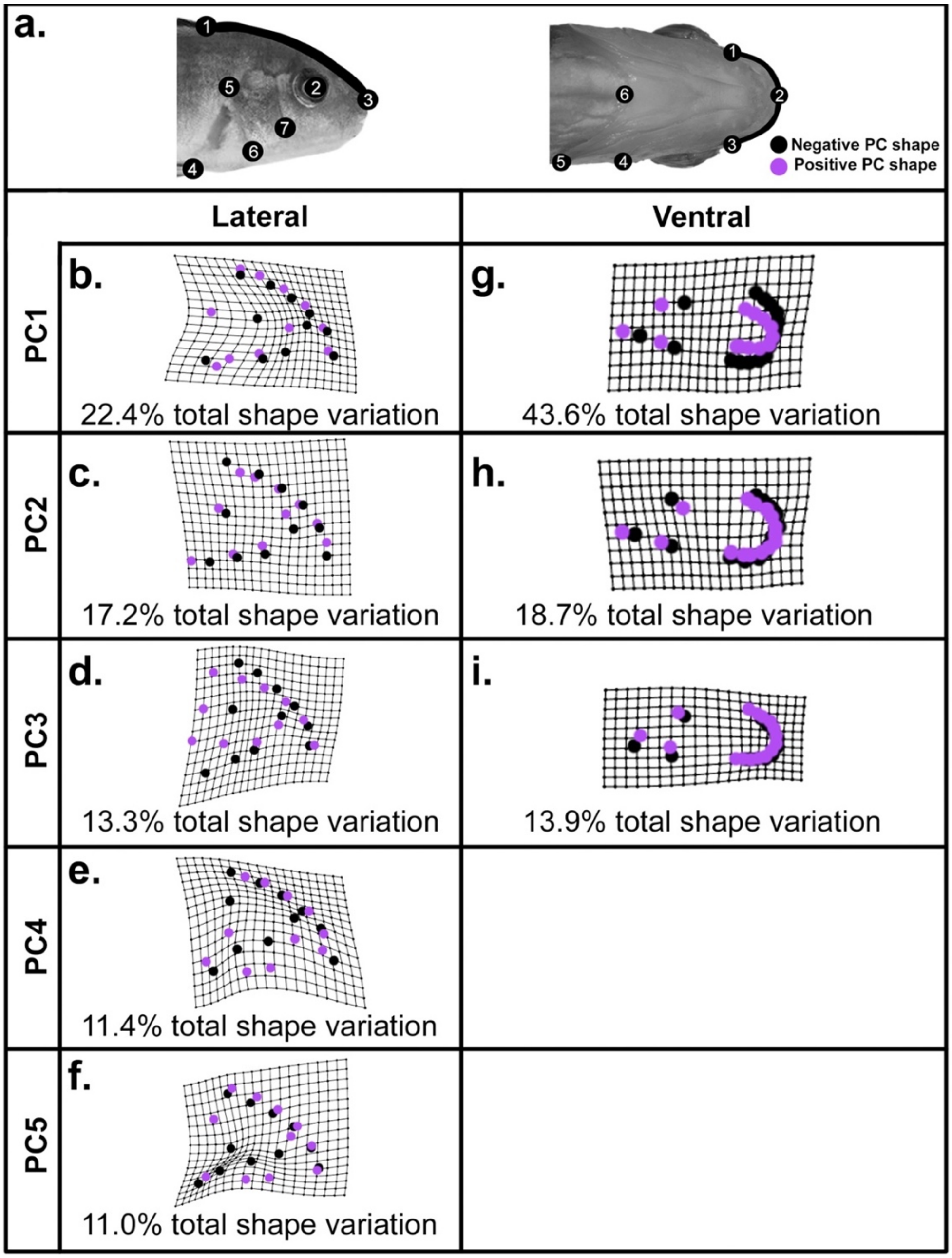


**Figure S1. Shapes indicated by PC scores following geometric morphometric analysis.** Landmarks used in analysis are as indicated in **(a)** and described in Figure 1. Shape illustrated by negative PC scores are in black dots and shapes for positive PC scores are in purple dots. Lateral shapes are in **(b-f)** and ventral analysis in **(g-i)** for **(b, g)** PC1 scores, **(c, h)** PC2 scores, **(d, i)** PC3 scores, **(e)** PC4 scores, and **(f)** PC5 scores. PC4 and PC5 scores did not contribute over 10% of shape variation and are thus omitted here. Total phenotypic variation explained by each PC is indicated.

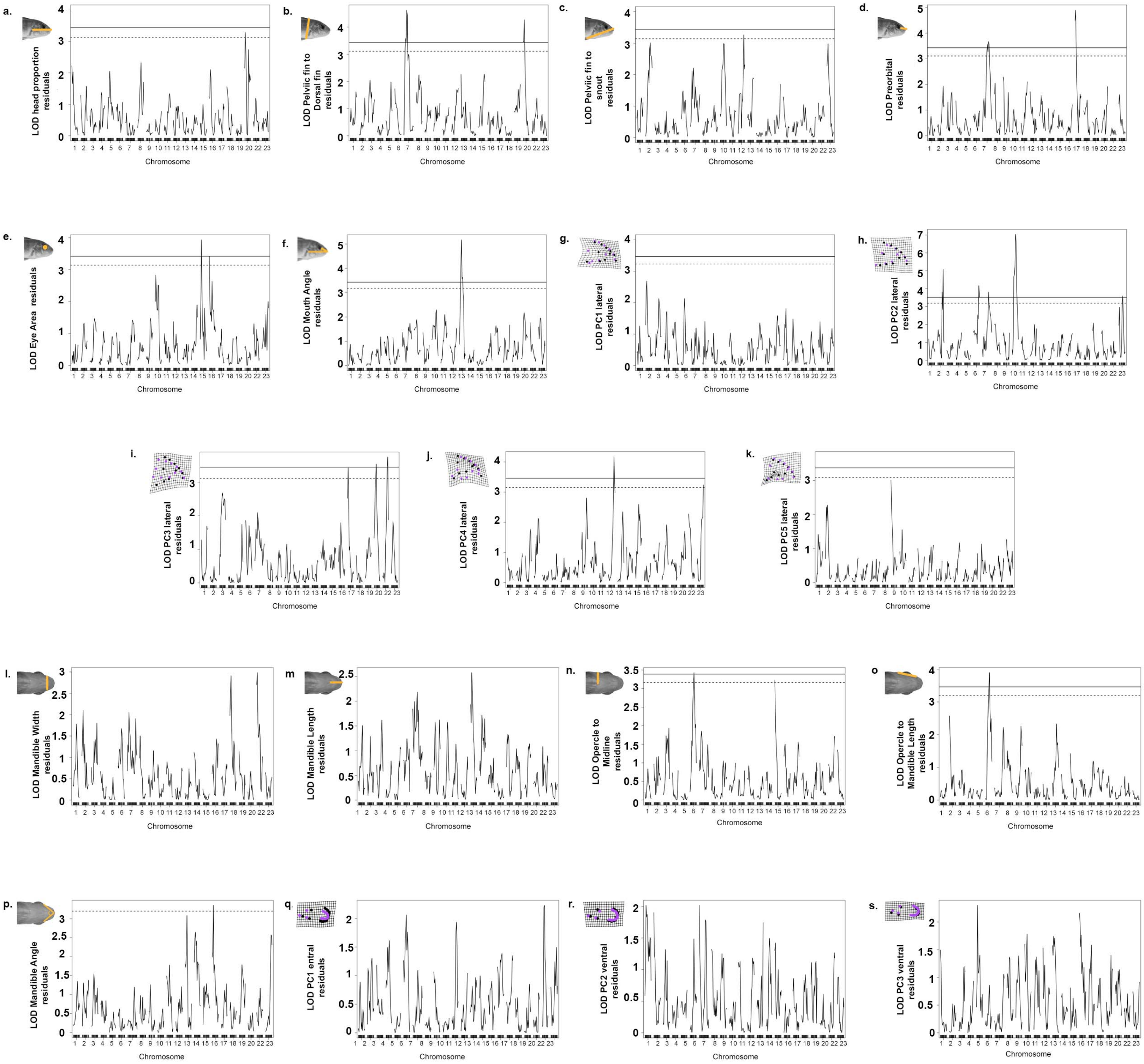


**Figure S2. Genome-wide QTL scans.** Phenotypic measures are indicated by illustration and include **(a)** head proportion, measured as head length/standard length, **(b)** dorsal to pelvic fin length, **(c)** snout to pelvic fin length, **(d)** length of the preorbital region of the head, **(e)** eye area, **(f)** mouth angle, **(g)** PC1 lateral shape, **(h)** PC2 lateral shape, **(i)** PC3 lateral shape, **(j)** PC4 lateral shape, **(k)** PC5 lateral shape, **(l)** mandible width, **(m)** mandible length, **(n)** opercle to midline width, **(o)** length from the opercle to the mandible, **(p)** angle formed from posterior ends of the mandible to the midline, **(q)** PC1 ventral shape, **(r)** PC2 ventral shape, and **(s)** PC3 ventral shape. Genome-wide significance is indicated at the 5% (solid line) and 10% (dashed line) level. No QTL meet significance if solid and dashed lines are not present. Details of QTL scans are in Table S3. QTL scans by chromosome are in Figure S3.

a.

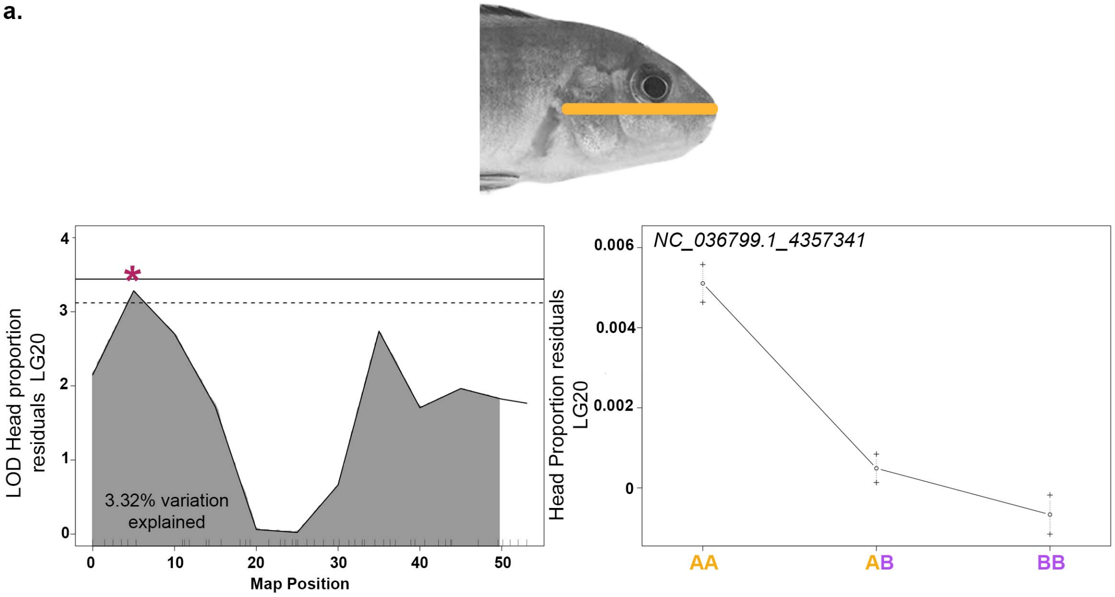

LOD Head proportion residuals LG20
3.32% variation explained
Map Position
NC_036799.1_4357341
Head Proportion residuals LG20
AA
AB
BB

**b.**

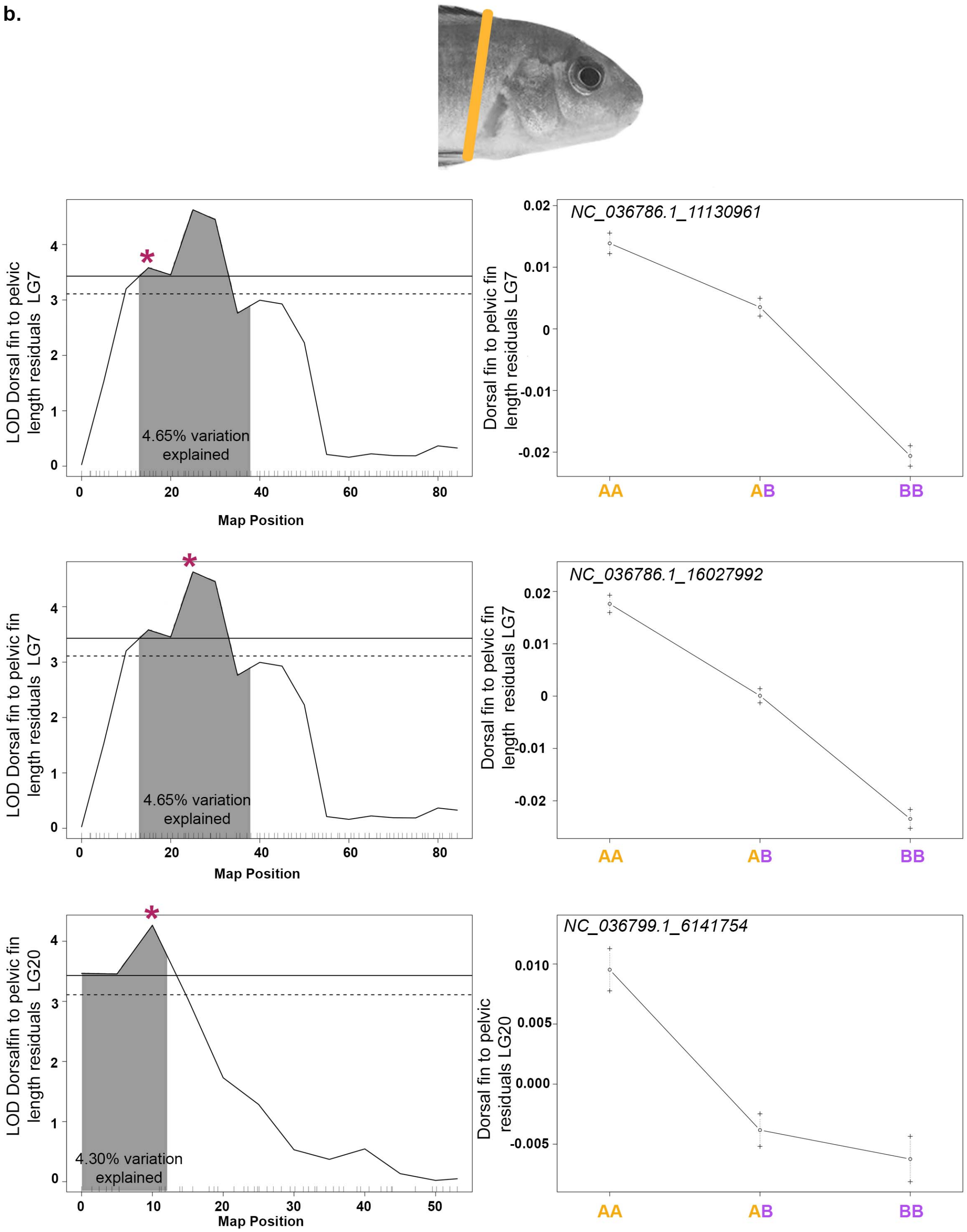

LOD Dorsal fin to pelvic fin length residuals LG7
4.65% variation explained
Map Position
NC_036786.1_11130961
Dorsal fin to pelvic fin length residuals LG7
AA
AB
BB
LOD Dorsal fin to pelvic fin length residuals LG7
4.65% variation explained
Map Position
NC_036786.1_16027992
Dorsal fin to pelvic fin length residuals LG7
AA
AB
BB
LOD Dorsalfin to pelvic fin length residuals LG20
4.30% variation explained
Map Position
NC_036799.1_6141754
Dorsal fin to pelvic residuals LG20
AA
AB
BB

**c.**

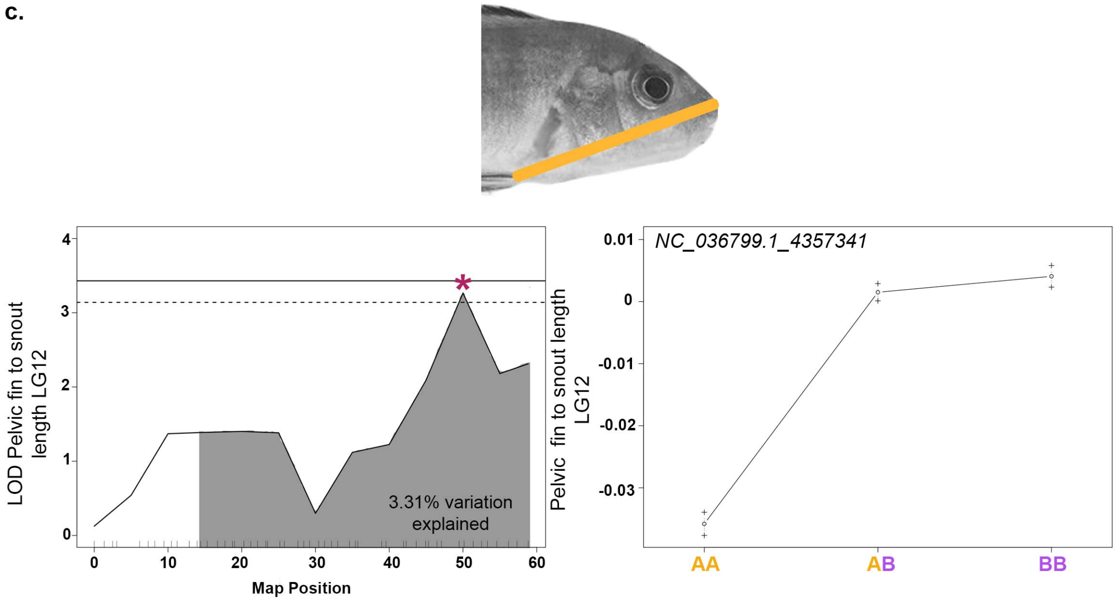

LOD Pelvic fin to snout length LG12
3.31% variation explained
Map Position
0
10
20
30
40
50
60
4
3
2
1
0
NC_036799.1_4357341
Pelvic fin to snout length LG12
0.01
0
-0.01
-0.02
-0.03
AA
AB
BB

**d.**

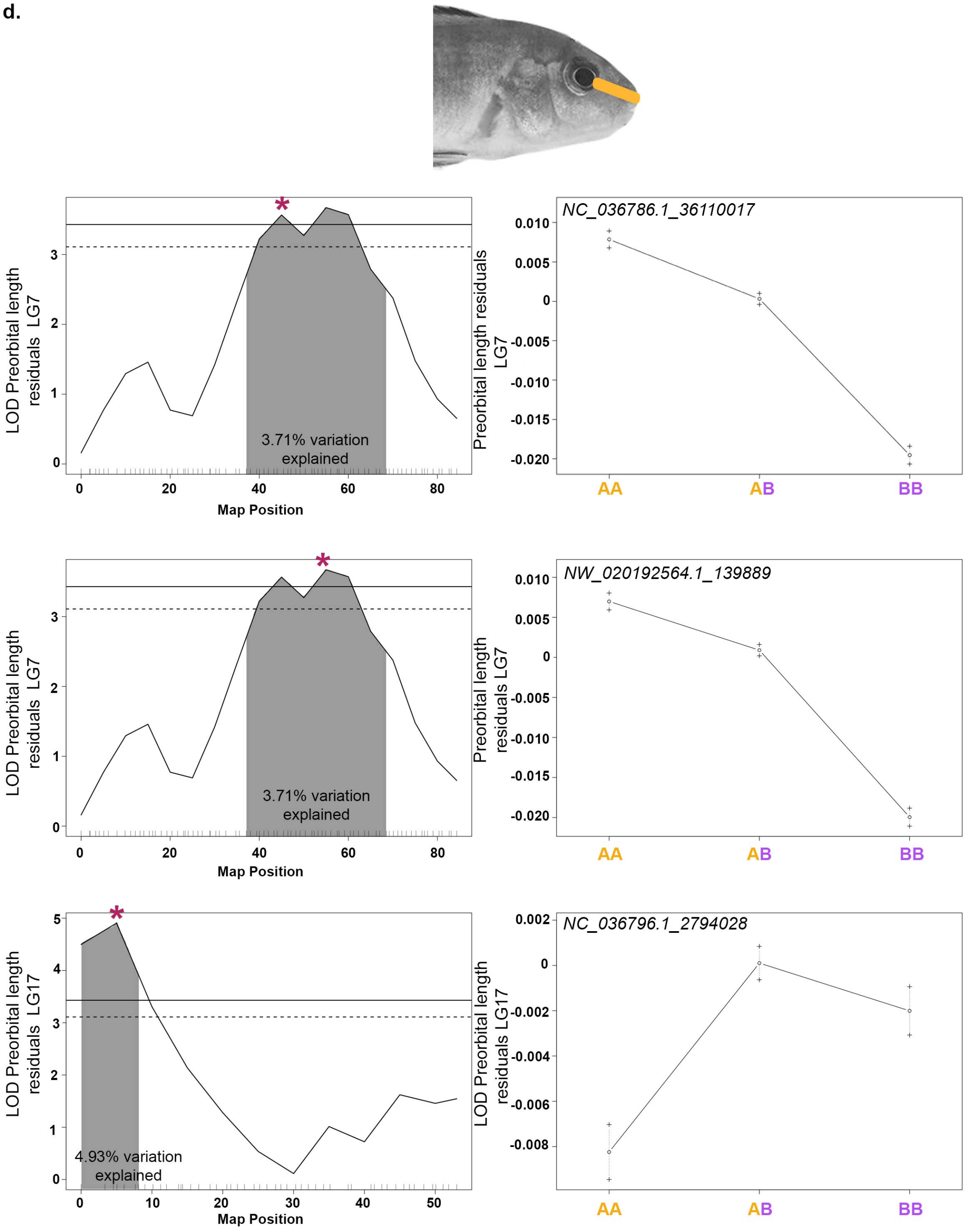

LOD Preorbital length residuals LG7
3.71% variation explained
Map Position
NC_036786.1_36110017
Preorbital length residuals LG7
AA
AB
BB
LOD Preorbital length residuals LG7
3.71% variation explained
Map Position
NW_020192564.1_139889
Preorbital length residuals LG7
AA
AB
BB
LOD Preorbital length residuals LG17
4.93% variation explained
Map Position
NC_036796.1_2794028
LOD Preorbital length residuals LG17
AA
AB
BB

**e.**

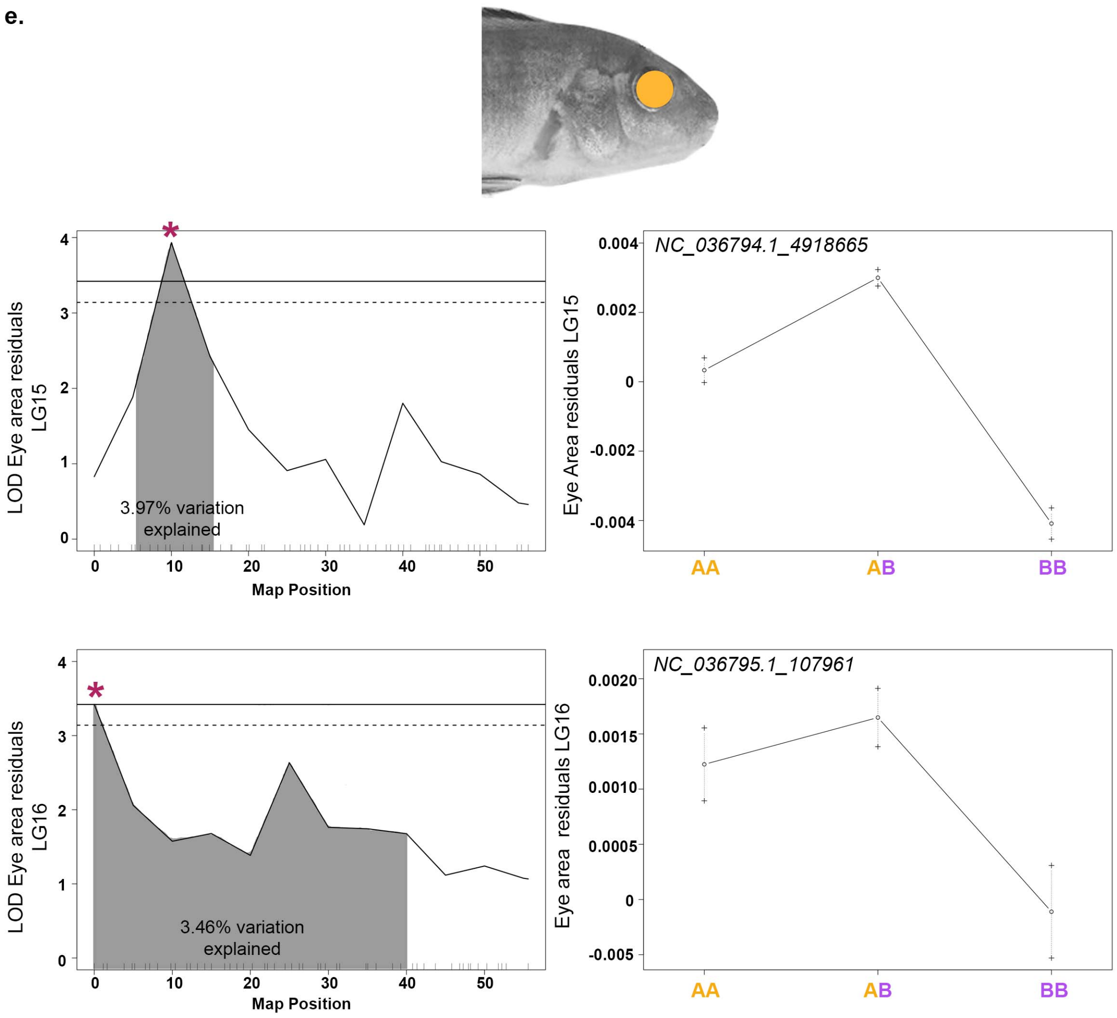
LOD Eye area residuals LG15
3.97% variation explained
Map Position
NC_036794.1_4918665
Eye Area residuals LG15
AA
AB
BB
LOD Eye area residuals LG16
3.46% variation explained
Map Position
NC_036795.1_107961
Eye area residuals LG16
AA
AB
BB

**f.**

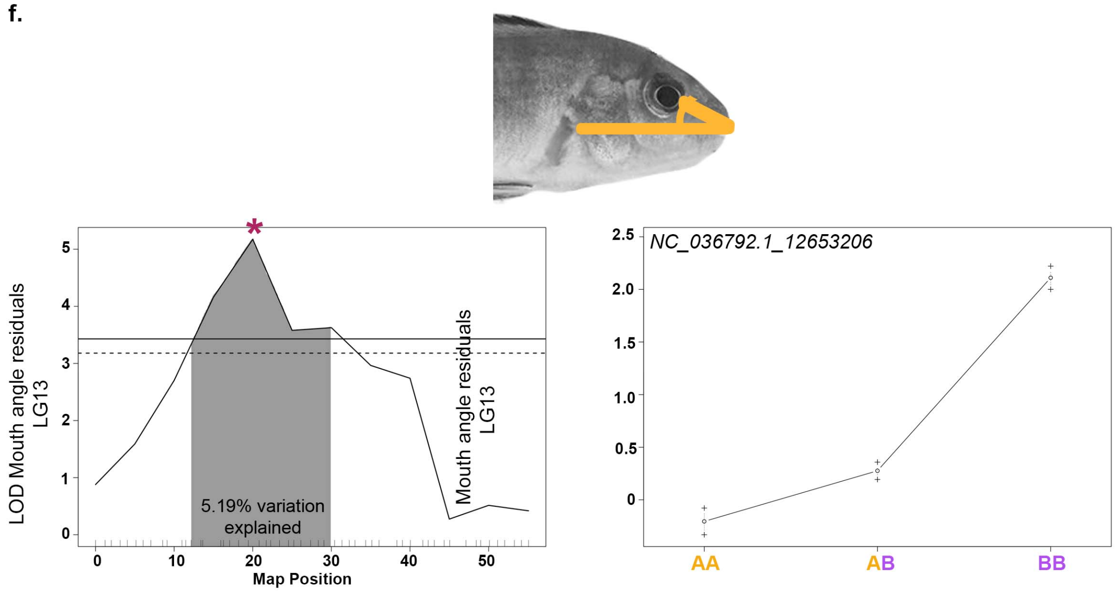
LOD Mouth angle residuals LG13
5.19% variation explained
Map Position
Mouth angle residuals LG13
NC_036792.1_12653206
AA
AB
BB

**g.**

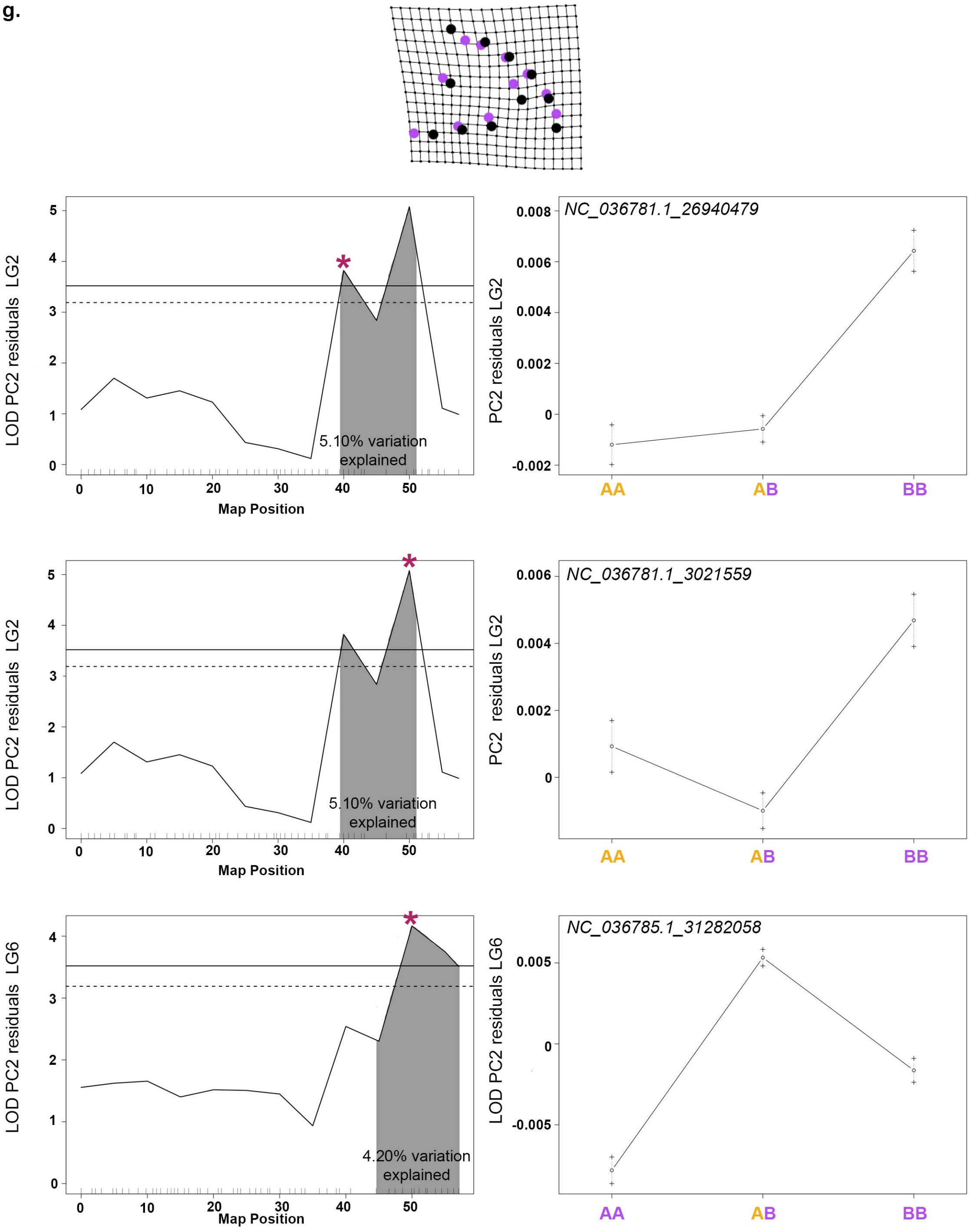

LOD PC2 residuals LG2
Map Position
5.10% variation explained
NC_036781.1_26940479
PC2 residuals LG2
AA
AB
BB
LOD PC2 residuals LG2
Map Position
5.10% variation explained
NC_036781.1_3021559
PC2 residuals LG2
AA
AB
BB
LOD PC2 residuals LG6
Map Position
4.20% variation explained
NC_036785.1_31282058
LOD PC2 residuals LG6
AA
AB
BB

h.

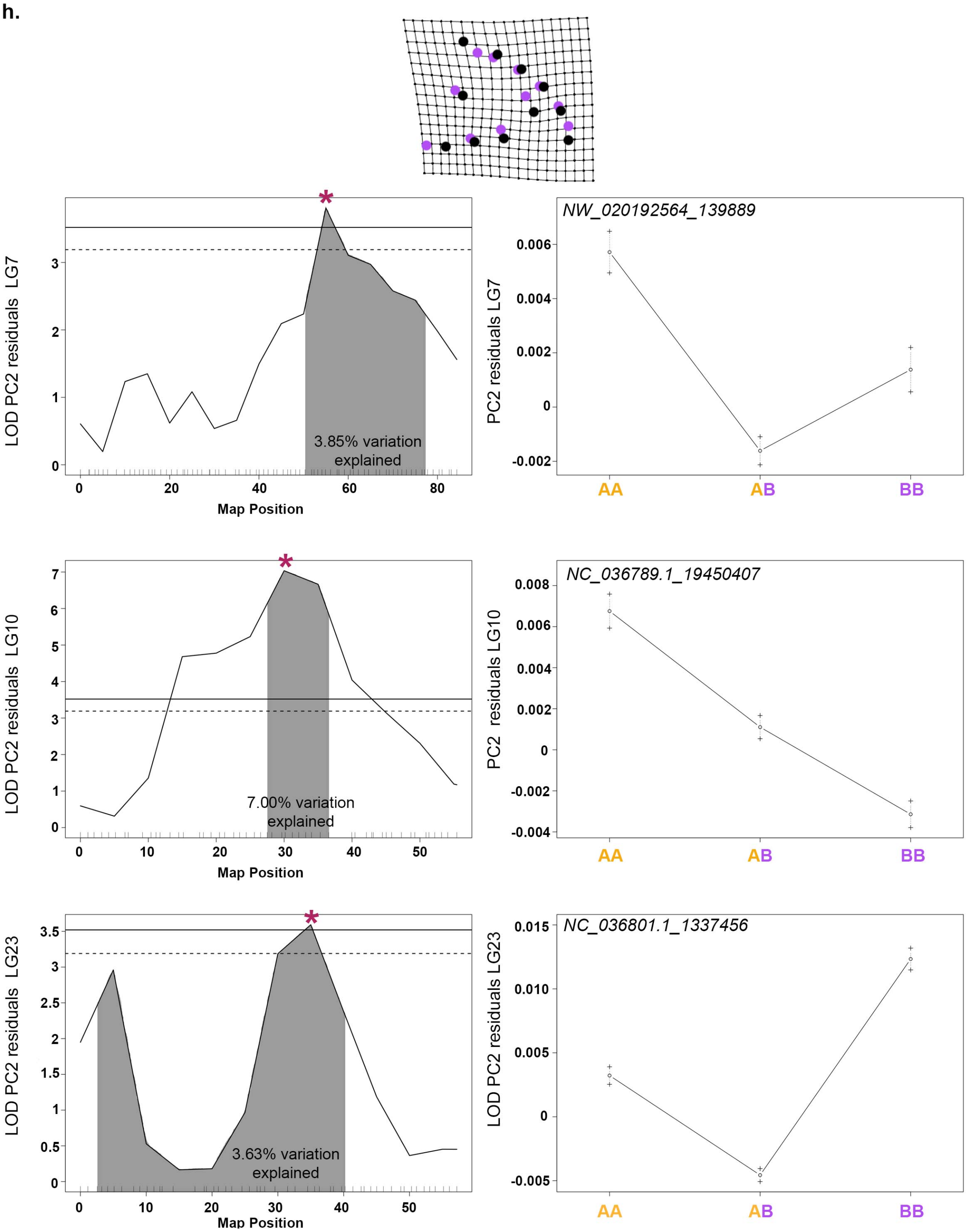

LOD PC2 residuals LG7
3.85% variation explained
Map Position
NW_020192564_139889
PC2 residuals LG7
AA
AB
BB
LOD PC2 residuals LG10
7.00% variation explained
Map Position
NC_036789.1_19450407
PC2 residuals LG10
AA
AB
BB
LOD PC2 residuals LG23
3.63% variation explained
Map Position
NC_036801.1_1337456
LOD PC2 residuals LG23
AA
AB
BB

i.

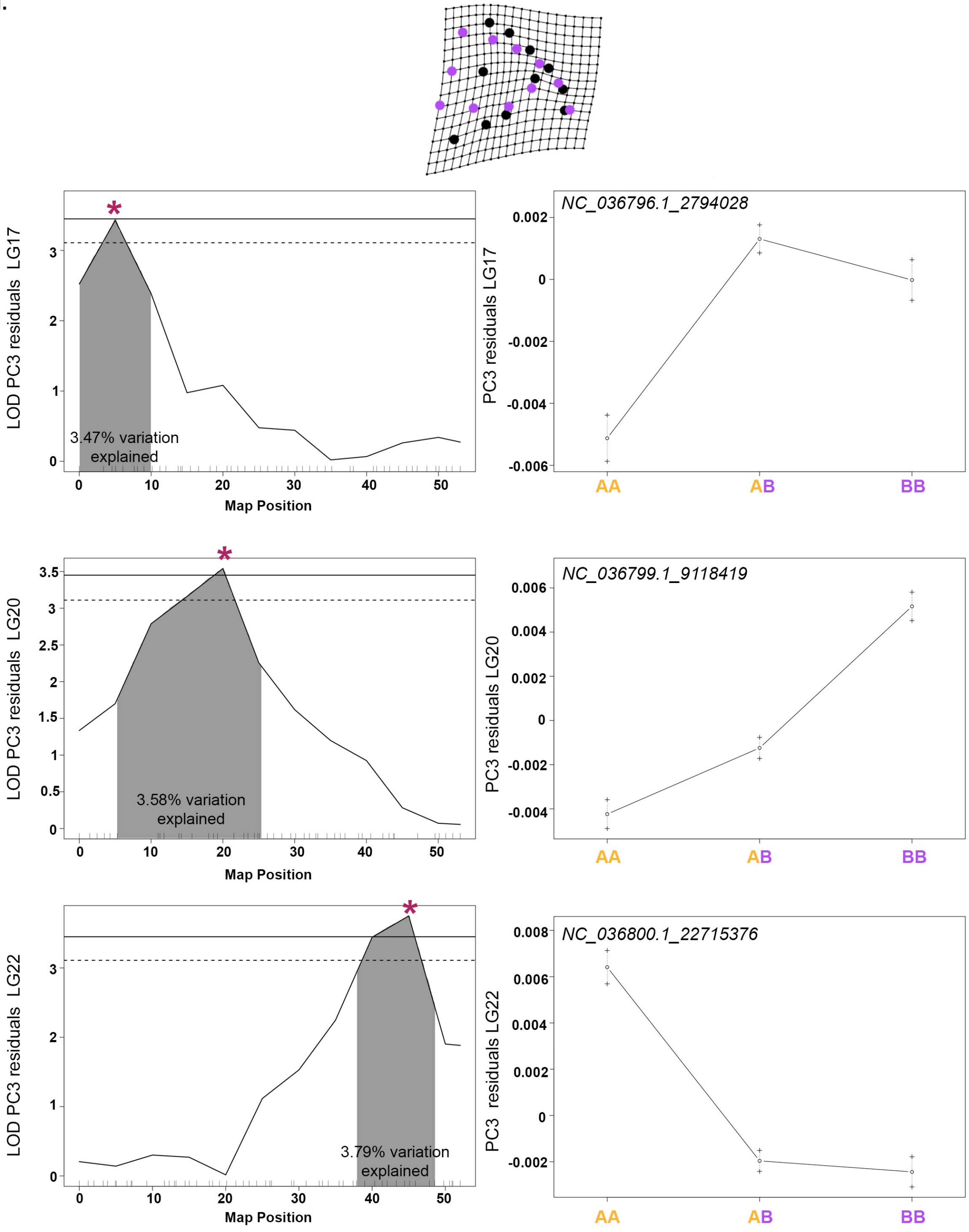

LOD PC3 residuals LG17
3.47% variation explained
Map Position
NC_036796.1_2794028
PC3 residuals LG17
AA
AB
BB
LOD PC3 residuals LG20
3.58% variation explained
Map Position
NC_036799.1_9118419
PC3 residuals LG20
AA
AB
BB
LOD PC3 residuals LG22
3.79% variation explained
Map Position
NC_036800.1_22715376
PC3 residuals LG22
AA
AB
BB

**j.**

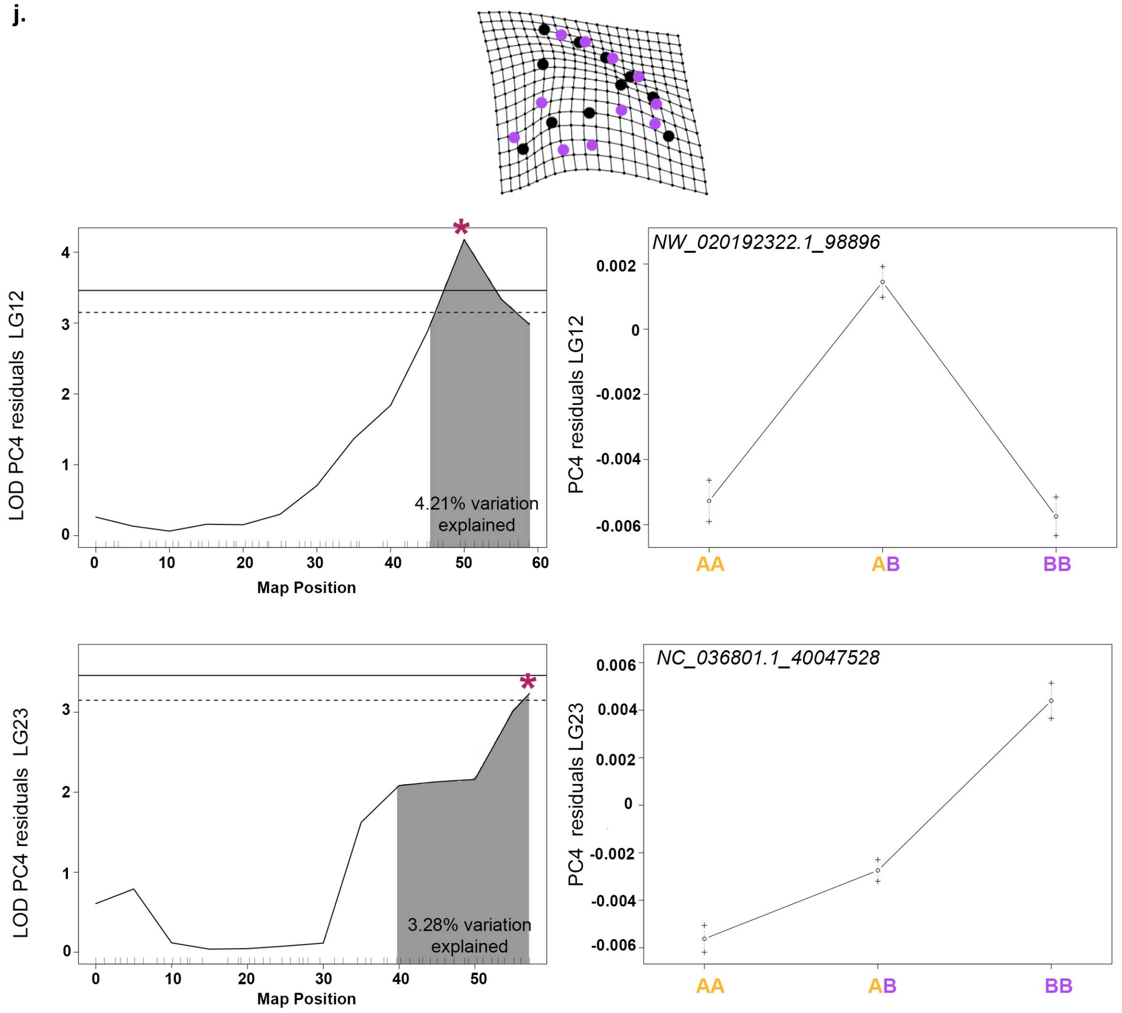

LOD PC4 residuals LG12
Map Position
4.21% variation explained
NW_020192322.1_98896
PC4 residuals LG12
AA
AB
BB
LOD PC4 residuals LG23
Map Position
3.28% variation explained
NC_036801.1_40047528
PC4 residuals LG23
AA
AB
BB

**k.**

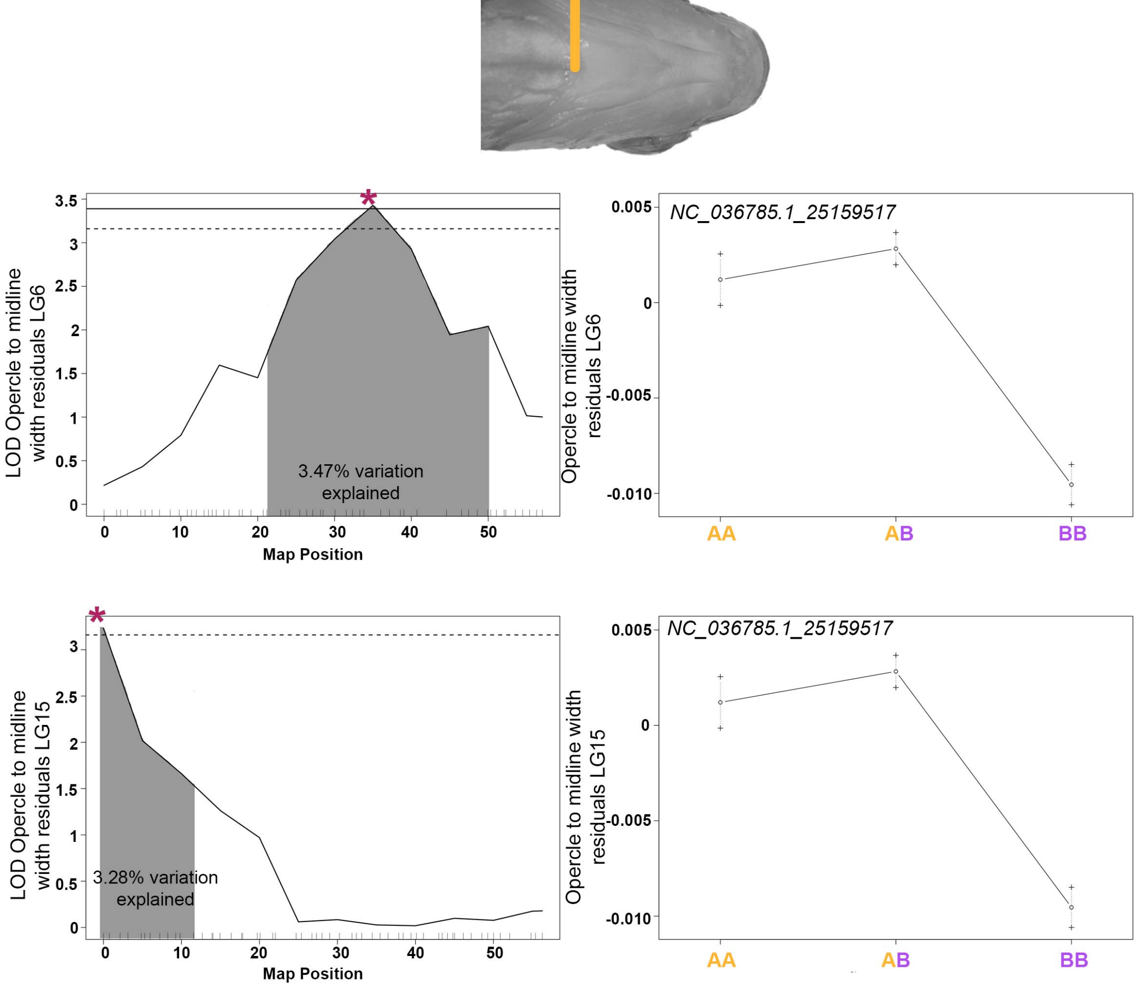

3.47% variation explained
LOD Opercle to midline width residuals LG6
Map Position
NC_036785.1_25159517
Opercle to midline width residuals LG6
AA
AB
BB
3.28% variation explained
LOD Opercle to midline width residuals LG15
Map Position
NC_036785.1_25159517
Opercle to midline width residuals LG15
AA
AB
BB

l.

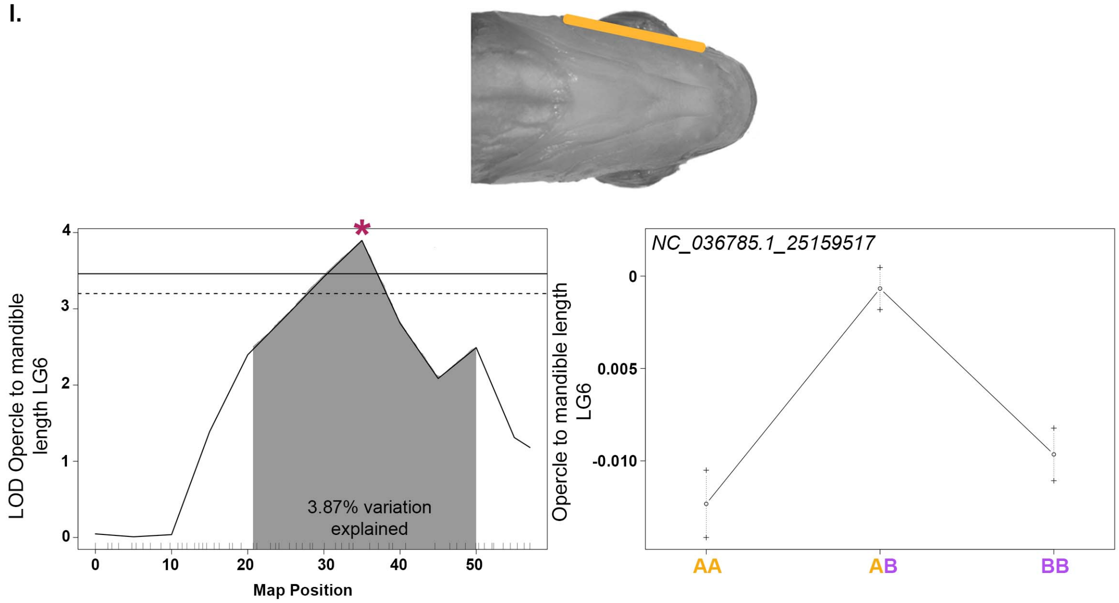


m.

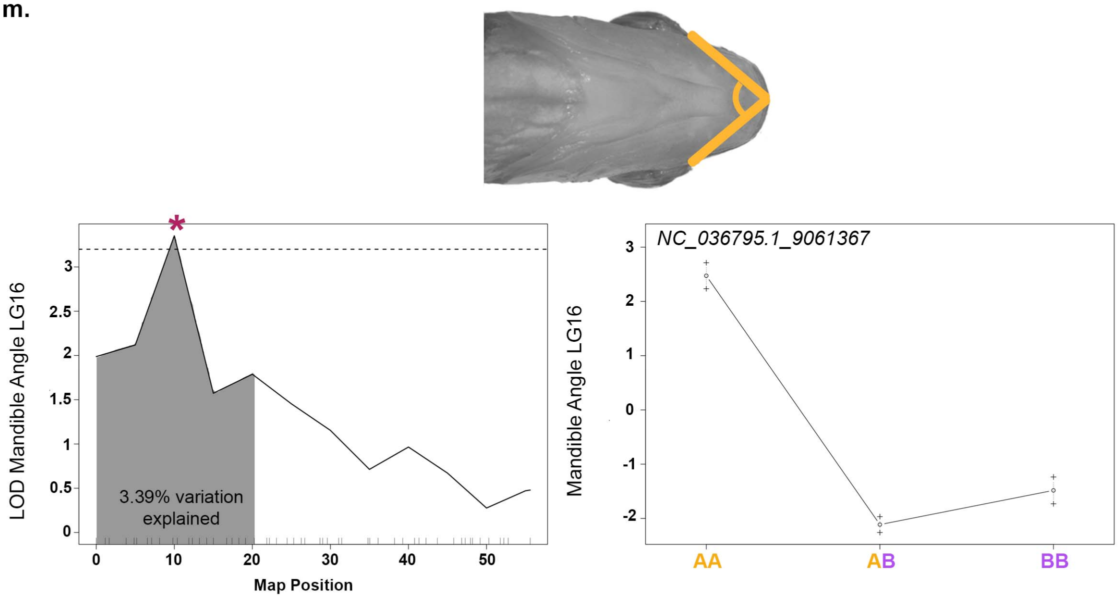


**Figure S3. QTL scans and allelic effects for QTL.** Phenotypic measures are indicated by illustration and include **(a)** head proportion, measured as head length/standard length, **(b)** dorsal to pelvic fin length, **(c)** snout to pelvic fin length, **(d)** length of the preorbital region of the head, **(e)** eye area, **(f)** mouth angle, **(g-h)** PC2 lateral shape, **(i)** PC3 lateral shape, **(j)** PC4 lateral shape, **(k)** opercle to midline width, **(l)** length from the opercle to the mandible, and **(m)** angle formed from posterior ends of the mandible to the midline. PC1 lateral shape, PC5 lateral shape, mandible width, mandible length, PC1 ventral shape, PC2 ventral shape, and PC3 ventral shape are not included in this figure as there were not any significant or suggestive QTL for these traits. 95% confidence interval for QTL is indicated by shading, percent of total phenotypic variation explained by QTL is reported, and genome-wide significance is shown at the 5% (solid line) and 10% (dashed line) level. Details of QTL scan are in Table S3 and genome-wide visuals are in Figure S2. Allelic effects are shown for the position indicated by *. The A allele was inherited from the *Labidochromis* granddam and the B allele from the *Labeotropheus* grandsire.
